# Transposon mutagenesis reveals differential essential pathways in model *Salmonella* Typhimurium strains SL1344 and SL3261

**DOI:** 10.1101/2023.10.03.556772

**Authors:** Jessica L. Rooke, Emily C. A. Goodall, Karthik Pullela, Rochelle Da Costa, Nicole Martinelli, Ian R. Henderson

## Abstract

*Salmonella enterica* is a globally disseminated pathogen that is the cause of over 100 million infections per year. The resulting diseases caused by *S. enterica* are dependent upon host susceptibility and the infecting serovar. For example, Typhoid fever is a human exclusive disease caused by *S. enterica* serovar Typhi. As *S. enterica* serovar Typhimurium induces a typhoid like disease in mice, this model has been used extensively to illuminate various aspects of *Salmonella* infection and host responses. However, the infection is so severe that even one infectious bacterium injected intravenously will cause mortality in 100% of animals within one week of infection. Due to this severity, researchers often use strains of mice resistant to infection or attenuated *Salmonella* strains to understand adaptive immunity and infection dynamics. Despite decades of research, many aspects of *Salmonella* infection and fundamental biology remain poorly understood. Here, we use a Transposon Insertion Sequencing (TIS) technique to interrogate the essential genomes of widely used isogenic wild-type and attenuated *S*. Typhimurium strains. We reveal differential essential pathways between strains, provide a direct link between iron starvation, DNA synthesis and bacterial membrane integrity, and show *S.* Typhi and *S.* Typhimurium have similar requirements for iron.

## Introduction

*Salmonella enterica* is a globally disseminated organism that can cause infections in both humans and animals. *S. enterica* is a facultative intracellular pathogen spread orofecally and is especially prevalent in areas that have poor sanitation or limited access to clean drinking water (1, 2). *S. enterica* can be broadly split into >2500 serovars, which differ in genome content and host restriction. There are multiple disease presentations of *S. enterica*, which depend upon the host sensitivity and infecting serovar (3). Some serovars are “generalists” that can infect many hosts (e.g. serovar Typhimurium), while others are specialists that can only infect a specific host, such as serovars Typhi and Paratyphi (4, 5).

*S. enterica* causes disease by actively invading epithelial cells of the intestinal mucosa via the well characterised SPI-1 type 3 secretion system (6). Invasive disease occurs when bacteria traverse the epithelial barrier and invade phagocytes at the basal surface of the gut lumen (7, 8). These phagocytes then circulate *Salmonella* to systemic sites through the lymphatic system (7, 9). The dynamics of *Salmonella* infections have largely been elucidated using a murine model of infection, using susceptible Nramp^-/-^ mouse strains that are incredibly susceptible to intracellular pathogens (10). Due to the severity of this disease model, to understand adaptive immunity and infection dynamics post day 7, researchers often use resistant mouse strains or attenuated *Salmonella* strains (11, 12). The infections caused in these models are less severe and are often cleared by the mice, and thus can be used to investigate later aspects of infection and immunity.

One popular attenuated strain is termed *S.* Typhimurium strain SL3261, which is an aromatic amino acid auxotroph developed by Hoiseth and Stocker via transposon mutagenesis (13). These researchers used a Tn*10* Tet^R^ transposon to mutate the parent strain *S.* Typhimurium strain SL1344. The corresponding transposon mutants were screened for tetracycline resistance and aromatic amino acid auxotrophy. The researchers then selected for aromatic amino acid auxotrophy and loss of tetracycline resistance, to generate SL3261. The resulting auxotrophy was due to transposon inactivation of the *aroA* gene in the shikimate pathway. Mutants in *aroA* are unable to synthesise chorismite, a molecule that sits at the branch point for many biosynthetic pathways including aromatic amino acid (13) and siderophore biosynthesis (14).

SL3261 was shown to be highly attenuated in a susceptible mouse model of infection and when used as a live attenuated vaccine candidate, provided protection against challenge with the virulent *S.* Typhimurium SL1344 strain (13). *S.* Typhimurium SL3261 has been a useful tool for investigating protective immunity towards *Salmonella* infections, bacterial disease burdens 7 days post infection, and is also used as a vector to deliver anti-cancer molecules into tumours (15). Despite this strain, and other similar strains, being used for decades in research, the exact mechanism for attenuation is unclear. Aromatic amino acids are abundant within the host (16) and so it is unlikely that loss of aromatic amino acid biosynthesis is the main driver of attenuation. While *aro* mutants have increased membrane permeability and susceptibility to membrane acting compounds, including human complement (17), the mechanism for this is also unknown.

Here we sought to understand the molecular mechanism for the attenuation of SL3261 to better understand the science in the field related to the use of this strain to elucidate *Salmonella* infection biology. We generated dense transposon mutant libraries in both *S*. Typhimurium SL3261 and the isogenic parent *S*. Typhimurium strain SL1344. Analyses of these libraries revealed major differences in genes involved in iron acquisition, DNA synthesis and cell envelope biogenesis. Importantly, these data indicate that SL3261 is restricted in ferric iron uptake, which is the primary cause of its membrane barrier defects and an attenuated phenotype.

## Results

### Construction of an *S.* Typhimurium strain SL1344 transposon library

TraDIS (Transposon Directed Insertion-site Sequencing) is a transposon insertion sequencing (TIS) method widely used to identify genes required for growth in specific conditions (18-24). To compare the genetic requirement of two *Salmonella* strains with near identical genotypes, but used in contrasting infection studies, we constructed two transposon-mutant libraries. *S*. Typhimurium transposon libraries have been previously constructed in strains SL1344 and SL3261 (25, 26) (Table S1). However, previously published SL1344 libraries were designed for screening in animal models and therefore had a low insertion density; where libraries of sufficient density were described, a catalogue of essential genes was not reported. As SL1344 is the parent strain of SL3261, we first constructed a library in *S*. Typhimurium strain SL1344. A library was constructed via transformation with an EZ-Tn*5* transposon and selected on agar plates supplemented with kanamycin (27, 28). Approximately 1.45 million individual colonies were pooled to form the library. To identify the transposon-insertion sites, genomic DNA (gDNA) was extracted from the library pool in duplicate and fragmented. DNA fragments containing the transposon-gDNA junction were amplified and sequenced using an Illumina MiSeq, obtaining >4 M reads per technical replicate. The data were de-multiplexed by barcode and searched for a correct transposon sequence. The barcodes and transposon sequences were trimmed, and the remaining data were mapped to the reference genome (accession NC_016810.1). Altogether, ∼10 M reads were successfully mapped for each library using the BioTradis analysis package (Table. S2). Some Tn*5* insertion-site preferences have been reported, however these are considered negligible in the context of an ultra-dense library (27, 29, 30). The transposon insertion sites are distributed around the whole chromosome and plasmids pSLT and pCol1B9 (Fig. 1A). However, we did not identify any transposon insertion events within the pRSF1010 plasmid (Fig. 1A), suggesting this plasmid may not be present in our strain or that it cannot tolerate insertions. The library replicates show high correlation (Fig. S1) and were pooled for subsequent analysis. Altogether, we identified 625,038 unique insertions in the chromosome and 20,545 and 22,017 unique insertions in the pCol1B9 (NC_017718.1) and pSLT (NC_017720.1) plasmids respectively. We identified genes required for growth using the BioTraDIS gene_essentiality.R script. Here, we excluded plasmids from our analysis firstly because *S.* Typhimurium SL1344 can be cured of plasmid pSLT (31) suggesting these genes are not required for viability, and secondly because of variability in plasmid copy number (between individual cells), this method of analysis can be inaccurate for identifying plasmid-genes required for fitness (32). We identified 516 chromosomal genes with significantly fewer transposon insertion events (Table. S3). A comparison between *S.* Typhimurium mutant libraries has been previously published (33), therefore we benchmarked our data against this dataset. Of the 516 essential genes in our dataset, 495 are protein coding and 363 of these (73.4%) are reportedly required for growth in all other *S*. Typhimurium strains (Fig. S2 and Table. S3). The proportion of genes unique to our library are consistent with equivalent datasets (31) and variation can be attributed to difference in methodology, conditions or analyses methods in addition to strain genotype discussed further below. This served as an important internal control for the validation of our library and a list of “core” *S.* Typhimurium essential genes are provided in Table S3.

**Figure 1.**
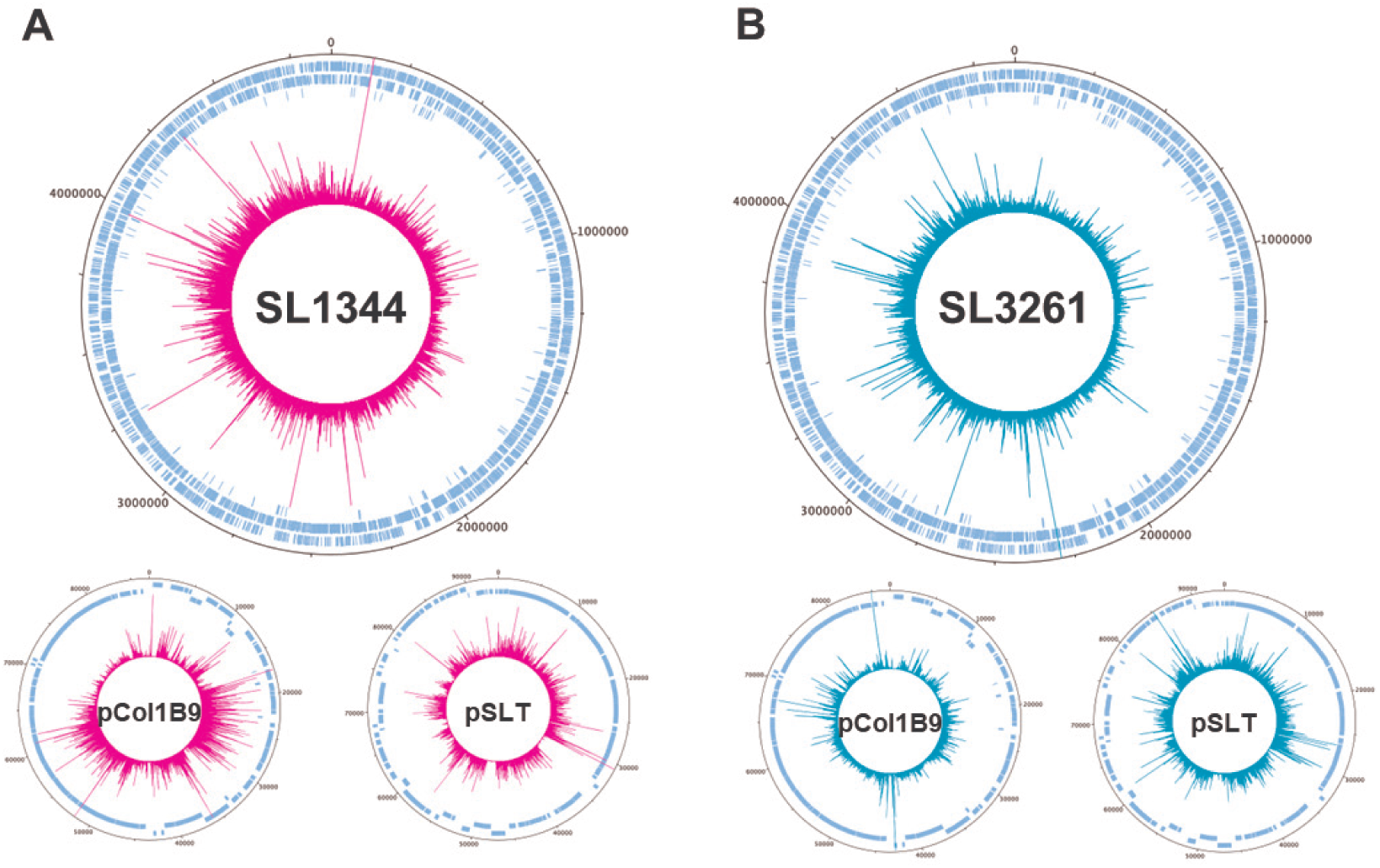
Transposon mutant libraries in *S.* Typhimurium strains. Chromosome and plasmid maps representing both transposon insertion site location and frequency in *S.* Typhimurium strains SL1344 (pink) and SL3261 (green).

### Construction of an *S.* Typhimurium strain SL3261 transposon library

Following the same protocol outlined above, we constructed a TraDIS library in S. Typhimurium strain SL3261. We identified 632,132 unique insertions for the chromosome, 23,431 for pCOL1B9 and 20,194 for pSLT. Like SL1344, we did not obtain any reads mapping to plasmid pRSF1010 (Fig. 1B) (Table. S2).

As previously noted, *aroA* encodes the enzyme 3-phosphoshikimate 1- carboxyvinyltransferase that is part of the shikimate pathway. This pathway generates chorismate, which sits as the branch point for many biosynthetic pathways including aromatic amino acids, ubiquinone, and siderophore biosynthesis (Fig. S3A). As expected, given SL3261 is an *aroA* mutant, no insertions were identified for the *aroA* locus (Fig. S3B). However, we also noticed that there were no insertions at the 5` end of the downstream *ycaL* gene. We hypothesised that this was possibly due to the way in which the SL3261 strain was constructed (13). Amplification of the *aroA-ycaL* loci (Fig. S3C) and subsequent Sanger sequencing confirmed the deletion of the *aroA* gene and revealed a scar comprised of an intact transposase gene that had replaced the 5′ end of *ycaL*, resulting in an in-frame truncation of *ycaL* (Fig. S3D). YcaL is a periplasmic M48 metalloprotease that is involved in outer membrane protein (OMP) biogenesis quality control that interacts with the β-barrel assembly (BAM) complex in *Escherichia coli* (34). M48 metalloproteases are conserved amongst the tree of life and are important for outer membrane integrity and OMP biogenesis (35). The truncation of *ycaL* in SL3261 resulted in the removal of the lipoprotein signal sequence (Fig. S3E), meaning if the protein was expressed, it would be mis-localised in the cytoplasm and not in the periplasm (Fig. S3F), essentially making SL3261 a double *aroA* and *ycaL* mutant. Therefore, we hypothesised the loss of YcaL may also contribute to SL3261 attenuation.

### Comparison of differential fitness between *S*. Typhimurium strains

To identify genes that differentially contribute to fitness in each strain, we used the biotradis tradis_comparison.R script. This method makes use of EdgeR to compare the relative abundance of sequencing read counts per gene for each library as a proxy for mutant abundance (36, 37). Mutants that are “more fit” are expected to be more abundant and therefore have a higher relative read count. Conversely mutants that are “less fit” will be less abundant and therefore have a lower relative read count. A comparison between the two strains revealed significant differences in genes involved in iron acquisition, DNA biosynthesis, outer membrane biogenesis and aromatic amino acid import (Fig. 2 and Table. S4). These data suggest that *S.* Typhimurium strain SL3261 has marked differences in cellular processes in comparison to *S*. Typhimurium strain SL1344 and these are explored further below.

**Figure 2.**
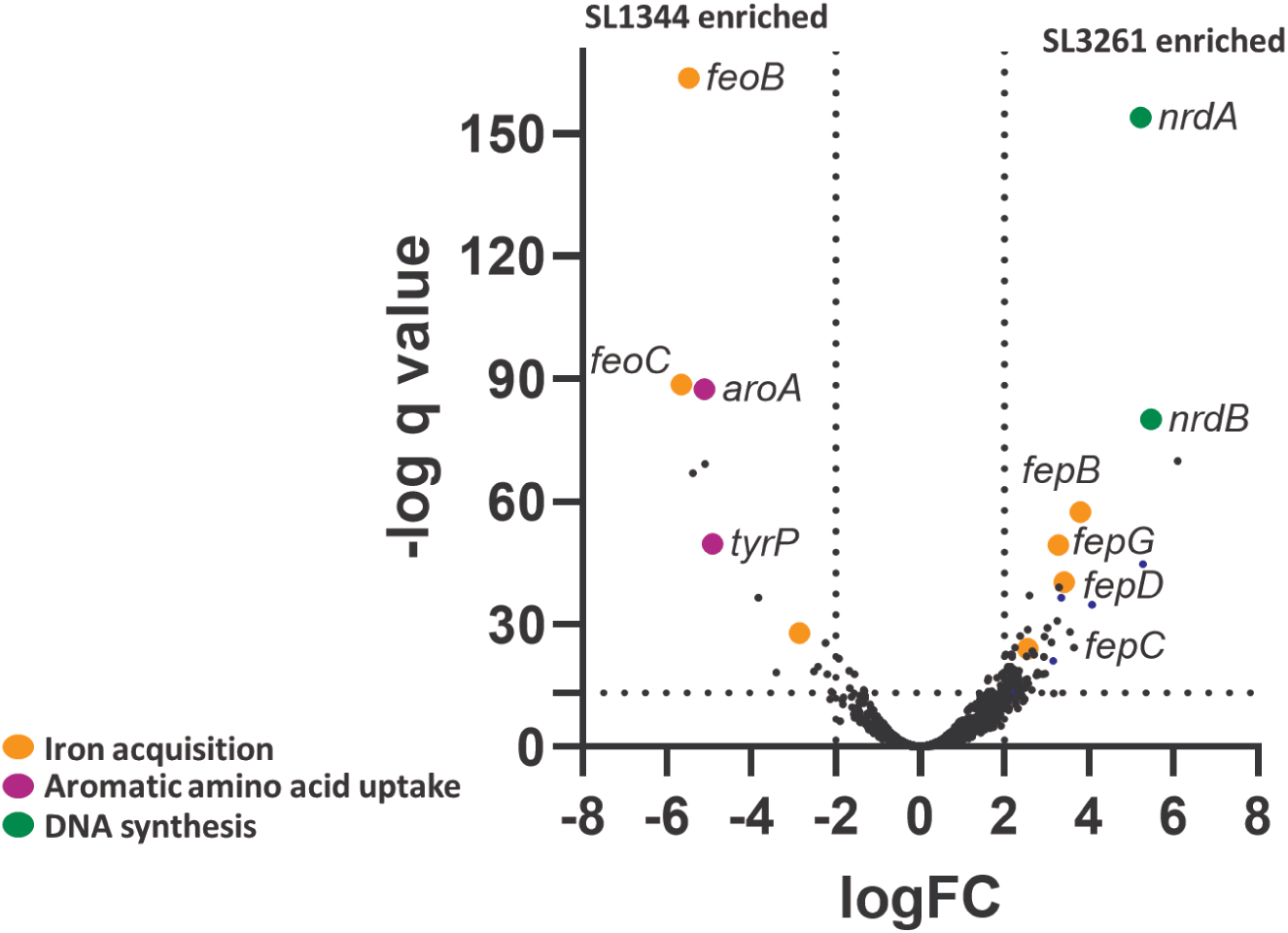
Identification of genes contributing to fitness in each strain. Read counts from each library were compared using edgeR to determine mutant fitness. The data are represented here as a volcano plot where mutants to the right of the y axis are enriched in SL3261, and the left are enriched in SL1344.

### Aromatic amino acid import

As SL3261 is unable to synthesise chorismate, a precursor to aromatic amino acid biosynthesis, we hypothesised that genes involved in import of aromatic amino acids would be essential. *S. enterica* and *E. coli* have four periplasmic aromatic amino acid permeases that transport aromatic amino acids through the cytoplasmic membrane, specific to each amino acid, using the proton motive force: L-tyrosine is imported by TyrP (38), L-phenylalanine by PheP (38) and L-tryptophan by Mtr (39) and the general aromatic amino acid permease AroP (40) (Fig. 3A). In our TIS data, the permeases for both phenylalanine and tryptophan were non-essential in both SL1344 and SL3261 whereas TyrP, a permease for tyrosine, was essential in SL3261 (Fig. 3B). AroP can transport all three aromatic amino acids (39), thus our TIS data would suggest one of the following: that under the conditions tested here AroP cannot, or is incredibly inefficient at transport of tyrosine; L-tyrosine is not as abundant as the other aromatic amino acids in LB agar and thus a specific L-tyrosine transporter is required; or the transporters are differentially regulated under laboratory conditions that result in only TyrP being essential in SL3261.

**Figure 3.**
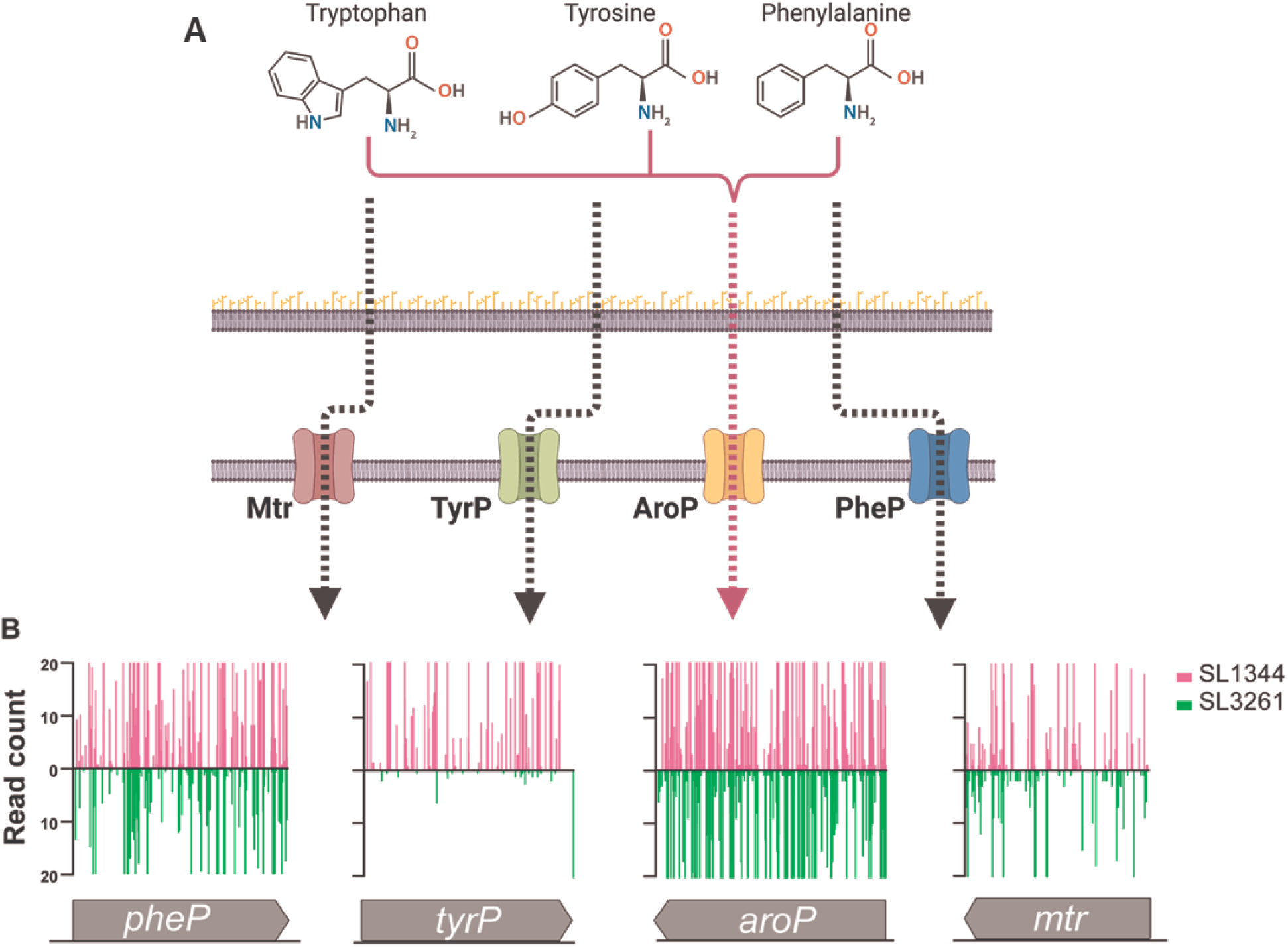
Aromatic amino acid transport mutant fitness. (B) Chemical structure of (left) phenylalanine, (middle) tyrosine and (right) tryptophan and their importers in the inner membrane (B) Gene insertion profiles of *pheP, tyrP, aroP* and *mtr* in SL1344 (pink) and Sl3261 (green) TIS libraries. For visualization, reads are capped at 20.

### Differential fitness of genes involved in iron utilisation

Iron is an important metal for biological life that is involved in various biological processes. Iron exists in two major forms: insoluble ferric (Fe^3+^) and soluble ferrous (Fe^2+^) forms, with ferrous iron considered the more unstable of the two, as it can readily undergo oxidation when exposed to environmental oxygen. The importance of iron import during *Salmonella* infections has been established previously (41-44) and mice that have impaired ability to remove iron from intracellular reservoirs are extremely susceptible to intracellular pathogens (10). The *S. enterica fep* and *feo* operons encode iron acquisition systems for the import of ferric and ferrous iron, respectively (45-47). Ferrous iron can enter the periplasm via outer membrane porins and is transported into the cytoplasm via the FeoAB-YhgG complex (45). In contrast, the import of ferric iron requires secreted siderophores that bind Fe^3+^ directly. The Fe^3+^- siderophore complex are imported through outer membrane porins such as FepA and IroN, a process that requires energy, which is derived from inner membrane complex TonB-ExbB-ExbD (48). Once in the periplasm, the iron-siderophore complex are dissociated and the iron transported from the periplasm to the cytosol via inner membrane transporter complex FepBCDG. *S. enterica* produces two siderophores: enterobactin and salmochelin.

The fitness of transposon mutants for these iron utilisation pathways were different between SL1344 and SL3261: SL1344 preferred import of ferric iron (*fepBCDG*) (Fig. 4A) whereas SL3261 preferred ferrous iron import (*feoAB yhgG*) (Fig. 4B). SL3261 was more sensitive to iron sequestration than SL1344 when we supplemented LB medium with the iron chelator 2-2-bipyridyl (2-BP) at multiple concentrations (Fig. 4C). The growth of SL3261 in the presence of 2-BP could not be restored by the supplementation of ferric iron into the culture medium, whereas topical addition of ferric iron restored growth of SL1344. These essentiality and growth differences are expected due to Aro mutants being unable to synthesise chorismate, which is a precursor for siderophores that are important for scavenging ferric iron from the environment. When we complemented SL3261 with *aroA* we restored the ability to withstand 2-BP mediated ferrous iron sequestration (Fig. S4A-D). However, what was intriguing was the apparent preference observed in iron utilisation systems in SL1344. We observed SL1344 *fep* mutants were less fit in or TIS data, and this can be observed in other TIS libraries in *S*. Typhi (26), iNTS strain D23580 (33) and *E. coli* K-12 (20). These data are intriguing, as presumably if the systems for siderophore mediated iron import are absent, strains can utilise the ferrous iron import system instead, and this might not cause a fitness disadvantage. However, our TIS data would suggest that mutants for the *fep* system are less fit *in vitro* and this is common amongst many *S. enterica* serovars.

**Figure 4.**
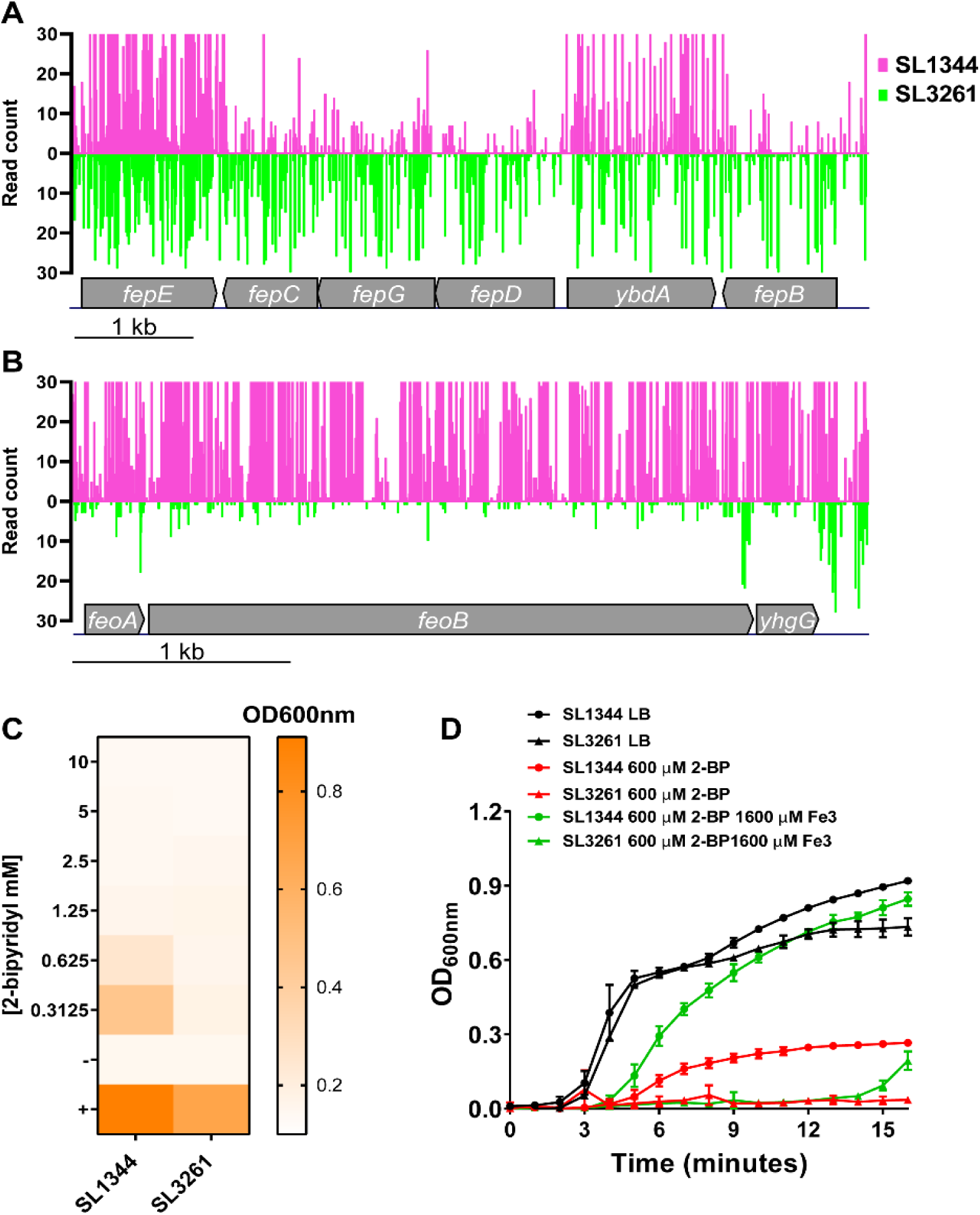
Ferric and ferrous iron importer fitness in SL1344 and SL3261. (A) Gene insertion profiles of the *fep* operon and (B) *feo* operon. (C) End point OD_600nm_ measurements for SL1344 and SL3261 grown in LB medium supplemented with varying concentrations of the iron chelator 2- bypyridyl (2-BP). (C) Growth curve of SL1344 and SL3261 in the presence of 2BP alone or 2-BP supplemented with Iron (III).

### Ribonucleotide reductase enzyme essentiality

The reduction of ribonucleotides (NTPs) to deoxyribonucleotides (dNTPs) is an essential process that occurs in all living organisms, that is achieved by ribonucleotide reductase (RNR) enzymes. There are three classes of RNR enzymes in *S. enterica*, which are differentially regulated (Fig. S5A) (49). The class Ia RNR operon encoded by *nrdAB* is the major RNR class expressed *in vitro* and is essential in *E. coli* and *S. enterica* (20, 26, 33, 50). The class Ib RNR operon is encoded by *nrdHIEF* and expression is induced by low iron conditions (51). The class III operon encoded by *nrdDG* is expressed under anaerobic conditions (52). In addition, each RNR enzyme complex requires a metal co-factor, and this is dependent upon the class (53) (Fig. S5A). NrdAB requires an Fe^3+^, whereas NrdHIEF requires Fe^3+^ or Mn^3+^ and NrdGD requires Fe^2+^. Our TIS data revealed that *nrdAB* is not essential is SL3261 (Fig. 5A). Surprisingly, when we interrogated our TIS data for the other RNR operons we observed that none are essential in SL3261 (Fig. 5B and C). A previous TIS study in *S*. Typhiurium suggested the essentiality of *nrdA* was iron dependent as *nrdA* transposon mutants were more fit in the presence of the iron chelator 2-dipyridyl (54, 55). In *E. coli*, *nrdAB* can be deleted provided there is an additional copy of *nrdHIEF* present either on the chromosome or a plasmid (56), suggesting that over expression of one of the other two RNR operons is sufficient to relive *nrdAB* essentiality. Due to the iron dependency of *nrdA* essentiality, differential RNR transcriptional regulation and iron co-factor requirement, we hypothesised that *nrdAB* transposon mutants become more fit in SL3261 due to ferric iron limitation. To overcome the lack of ferric iron and the need for RNR enzyme function, we hypothesised that the expression of either *nrdHIEF* or *nrdGD* would be increased in SL3261 compared to SL1344. To test this hypothesis, we generated luciferase reporter constructs whereby luminescence expression is driven by *nrd* promoters in the vector pLUX as well as a control promoter *gyrA.* We tested the luminescence intensity of each construct in *S*. Typhimurium SL1344 under three conditions: mid-exponential phase (MEP), anaerobic shock and low iron (Fe) shock (49). We show that the *nrdA* promoter shows similar activity in all conditions tested whereas the *nrdD* promoter was significantly more active under anaerobic conditions and *nrdH* under low iron (Fig. S5B), as expected. These data demonstrate that the reporter constructs respond appropriately in SL1344. To understand whether the *nrd* genes are regulated differently in SL3261, we transformed our reporter plasmids into SL3261 and analysed the relative luminescence intensity generated by each construct compared to the *gyrA* promoter over 16 hours of growth on LB agar, replicating selection conditions of our TIS library construction. We observed that there were no significant differences of either the *nrdA* and *nrdD* promoters between SL1344 and SL3261, but we did observe a different pattern of *nrdH* promoter activity between strains (Fig. 5D). These data suggest that the RNR operons are differentially regulated between strains and that in SL3261, NrdHIEF is expressed at higher levels than in SL1344 at certain points during growth and this can explain the apparent fitness of NrdAB transposon mutants in this background.

**Figure 5.**
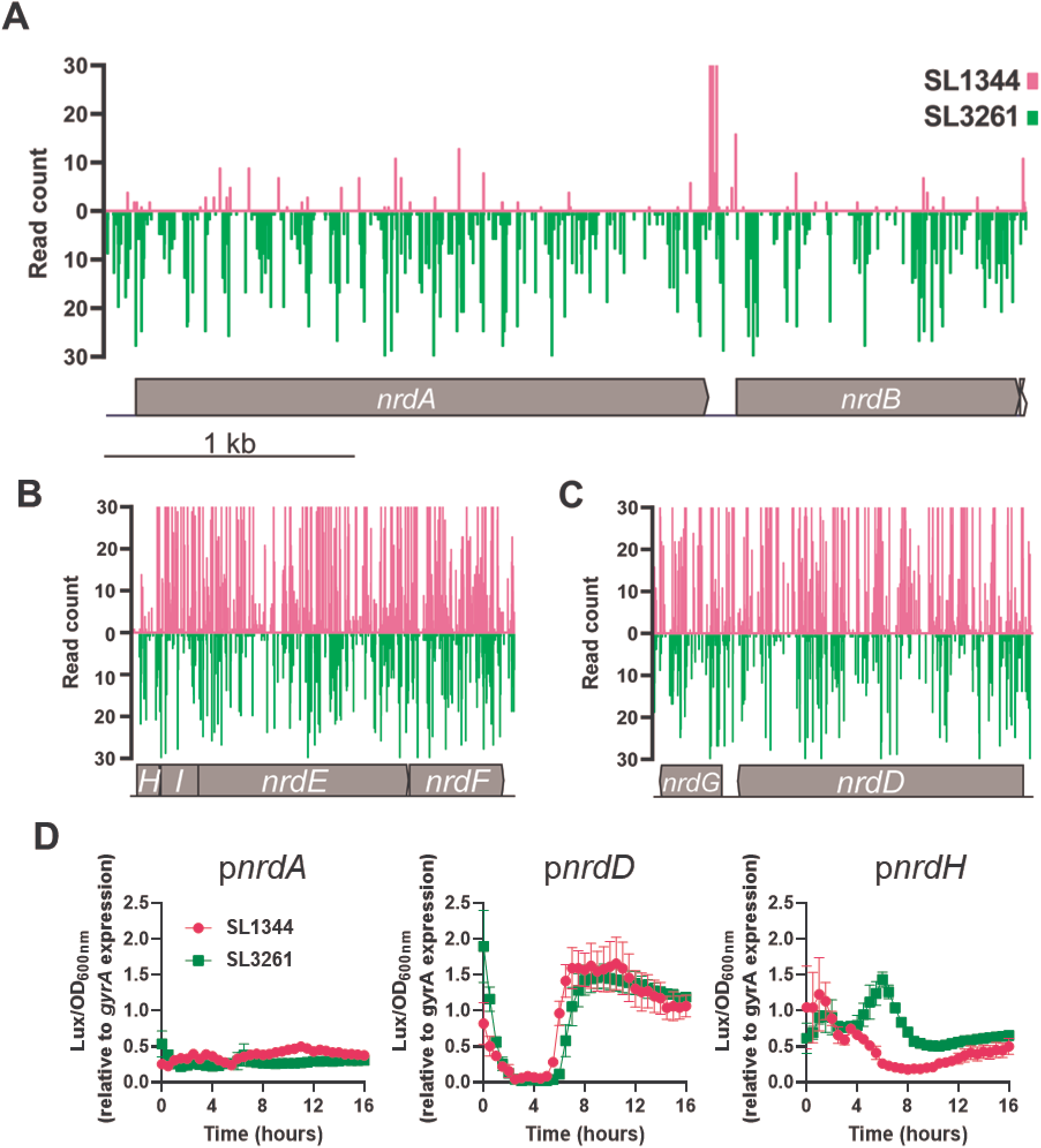
The role of ribonucleotide reductases in the absence of *aroA*. Insertion profiles of (A) *nrdAB*, (B) *nrdHIEF* and (C) *nrdGD* in the SL1344 (pink) and SL3261 (green) TIS libraries. (D) Relative expression of *nrd* genes compared to the *gyrA* promoter over 16 h of growth on LB agar. Data here from 3 biological replicates and error bars showing standard error of the mean (SEM).

### Fitness of genes involved in outer membrane biogenesis

The Gram-negative bacterial outer membrane is asymmetric and comprised of phospholipids at the inner leaflet and lipopolysaccharide (LPS) in the outer leaflet. LPS is an activator of host innate immunity via toll like receptor 4 (TLR4) signalling (57) and is important for barrier function of the outer membrane (58). In *Salmonella enterica*, LPS is comprised of lipid A, inner and outer core oligosaccharide, and O-antigen. In *S. enterica* group B (O4) serotypes, such as SL1344 and SL3261, the O-antigen is composed of repeating units of galactose-rhamnose-mannose-abequose sugars (59) (Fig. S6A). The addition of sugar subunits to core oligosaccharide and O-antigen are co-ordinated by multiple enzymes that are arranged in multiple operons (59). Surprisingly, mutations in several genes involved in LPS and O-antigen biosynthesis conferred increased fitness on SL3261 compared to SL1344 (Fig. S6A-D), despite no obvious differences when we compared the LPS profiles of both SL1344 and SL3261 (Fig S6E). Some of these gene clusters were also identified in a screen for transposon mutants that were differentially fit under iron restricted conditions (60), suggesting that the differential fitness we observe in our dataset can also be attributed to limited iron availability in SL3261.

Aro mutants are known to have outer membrane permeability defects and to be susceptible to compounds that are membrane acting, such as EDTA, albumin, complement and bile salts (17). Given YcaL is purported to interact with the Bam complex during OMP biogenesis, we hypothesised that loss of YcaL in SL3261 may disturb the OM barrier function. To test this hypothesis, we screened our strains and relevant complements on membrane acting compounds: vancomycin, bile salts and EDTA. We observed that for each compound, complementation of AroA restored membrane barrier permeability but complementation with YcaL alone did not (Fig. 6A). As *aro* mutants are deficient in synthesising aromatic amino acids and siderophores, we wanted to uncouple the mechanism behind the membrane barrier defect observed.

**Figure 6.**
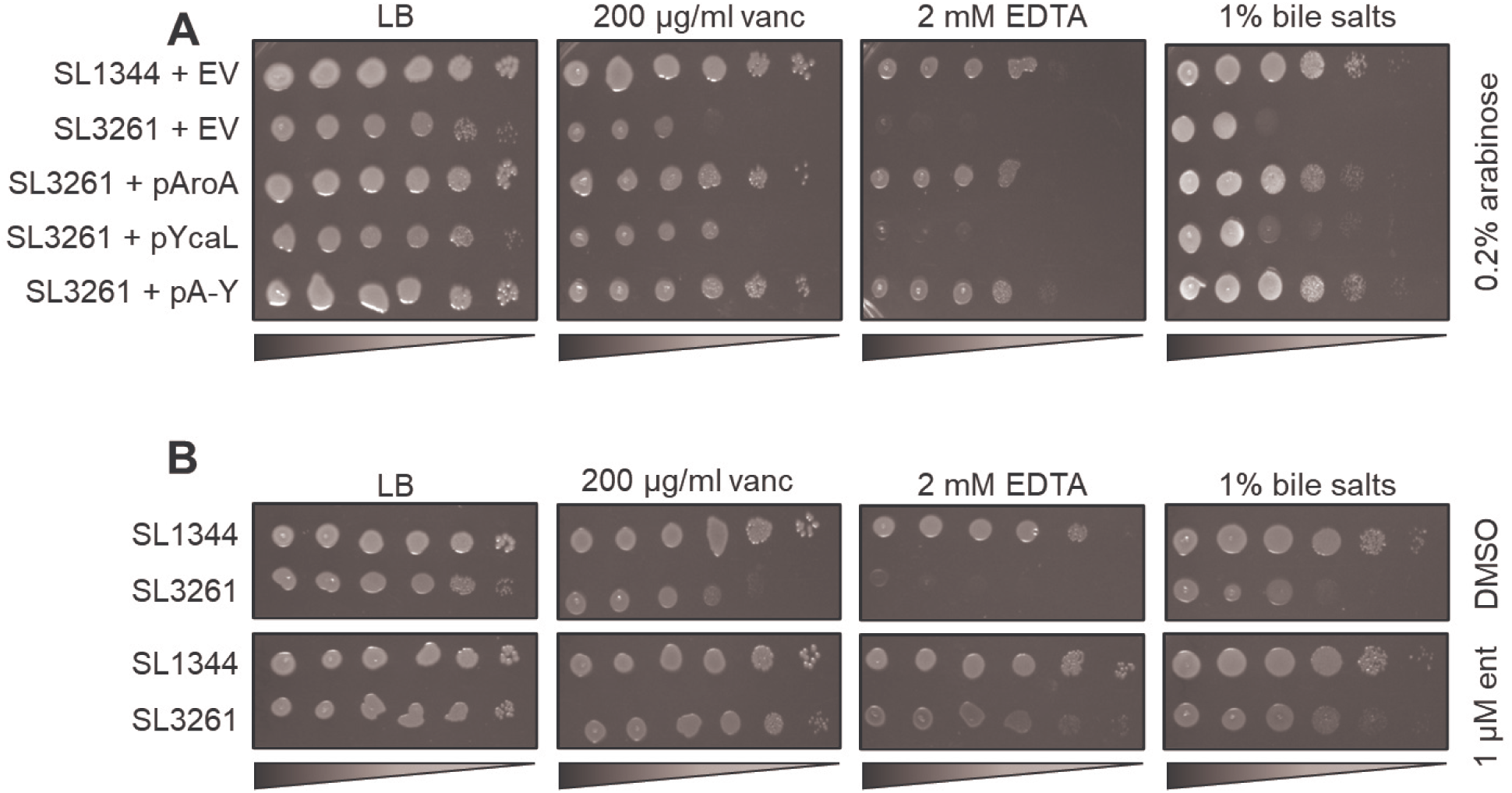
Outer membrane permeability screens for SL1344 and SL3261. (A) Overnight cultures of each strain were serially diluted, and spot plated onto LB agar supplemented with 0.2% arabinose and either 200 µg/ml vancomycin, 2 mM EDTA or 1% bile salts. (B) Overnight strains serially diluted and plated onto LB agar plates supplemented with the same membrane acting compounds and either DMSO or 1 µm enterobactin. Plates were incubated at 37°C for 16-18 h.

To that end, we repeated our chemical screens with SL1344 and SL3261 and either supplemented the agar with or without 1 µM enterobactin. We observed that enterobactin completely restored barrier function in SL3261 (Fig. 6B), suggesting iron limitation causes the barrier defects observed in SL3261.

## Discussion

TIS methodologies have been used to identified essential genomes of many bacterial pathogens (18, 20, 23, 25, 26, 33). Here, we compared the essential gene list of the extensively used laboratory *S*. Typhimurium strain SL1344 with other Typhimurium TIS datasets (33). We revealed a list of 363 Typhimurium genes that are essential in all strains tested. Differences in essential genes can arise due to differences in how libraries were constructed, and the data analysed. For example, our libraries were constructed without salt to increase transformation efficiency, and this might explain some of the differences in our essential gene lists compared to other published datasets.

Here we used TIS to compare transposon mutant fitness between two model strains of *S*. Typhimurium: SL1344 and SL3261. One of the main findings of this study was that transposon mutants responsible for iron utilisation were differentially fit between strains and that iron limitation heavily influenced SL3261 phenotypes. We observed that *S*. Typhimurium SL1344 preferred ferric iron whereas SL3261 preferred ferrous iron, due to an inability to synthesise siderophores. This apparent preference for ferric iron appears to be conserved amongst other *Salmonella* serovars, including Typhi. In fact, a previous study concluded that serovars Typhi and Typhimurium prefer utilising different iron: Typhi with ferric and Typhimurium ferrous, and that this was due to the different niches that these serovars occupy during human infection (26). However, this study did not account for the usage of SL3261 as the Typhimurium strain, and as we have shown here, this strain cannot use ferric iron due to the abolition of the siderophore biosynthetic pathway. Our study, and others, have now demonstrated that many serovars of *Salmonella* prefer ferric iron and this is not limited to Typhi, suggesting that iron utilisation does not differ between serovars or infectious niches.

SL3261 is highly attenuated in mice, and *aroA* mutants have been identified as attenuating in multiple serovars of *Salmonella* and other bacterial pathogens (61-64). These data suggested that synthesising aromatic amino acids was important *in vivo* and that perhaps these amino acids were in short supply in the host. However, recent data suggests that aromatic amino acids are not restricted inside the host and that mutants in genes that are dedicated to making aromatic amino acids are not attenuated (25). Given that *aroA* mutants are unable to synthesise chorismate, which sits at the branchpoint for at least three different biosynthetic pathways, these data suggest that the cause of attenuation in *aro* mutants was likely due to functions other than aromatic amino acid auxotrophy. In addition, *aroA* was determined to be important for *S*. Typhimurium colonisation on alfalfa sprout roots, and this attenuation could not be rescued by topical addition of neither tryptophan nor phenylalanine (65). This study also demonstrated that topical addition of ferrous iron could restore root colonisation and it was concluded that the lack of siderophore production and not lack of aromatic amino acids was important for *S*. Typhimurium colonisation of alfalfa roots (65).

Genes involved in ferric iron import are known to be important for infections (43) and the *fep* operon is upregulated inside macrophages and the cytosol of host cells (42, 49). There is also evidence that *nrdHIEF* is upregulated in host cells, suggesting iron limitation and that the major RNR enzymes used to synthesise DNA are not NrdAB during infections and therefore different to standard laboratory conditions (42, 49, 66). We also know that susceptible mouse strains lacking a functional copy of the protein Nramp1 are incredibly susceptible to intracellular infections, including *Salmonella* (10). *Nramp1* encodes a divalent cation transporter that can remove iron from host cell vacuoles and stimulate lipochalin-2 expression, a host iron chelator, thereby reducing intracellular iron concentrations (67, 68). In the absence of Nramp1, murine macrophages are unable to limit intracellular iron as efficiently, and therefore these macrophages are more susceptible to infection. Mouse strains considered resistant to *S. enterica* infections have a functional copy of Nramp1, and therefore the reason for the resistance to *Salmonella* infections is because these mice can efficiently restrict intracellular iron concentrations.

Gene mutations that cause membrane barrier defects can cause *S*. Typhimurium to become susceptible to membrane targeting compounds such as cationic antimicrobial peptides and detergents, such as bile salts, resulting in attenuation during infection (69). Aro mutants are known to have membrane barrier defects and here we show that barrier function can be restored in SL3261 by complementation of AroA on a plasmid and supplementation of enterobactin. As enterobactin supplementation completely restored barrier function in SL3261, we stipulate that in addition to iron limitation, SL3261 would experience severe membrane defects *in vivo*. This additional attenuating phenotype would contribute to the severe phenotype observed during SL3261 infections.

During this study we identified that SL3261 is essentially a double mutant of *ycaL* and *aroA*. YcaL is a metalloprotease that interacts with the Bam complex to facilitate OMP folding (34). Here, we could not identify any mutant fitness defects that could be obviously, or experimentally, attributed to the loss of YcaL. High throughput screens have also failed to find an obvious phenotype for YcaL in *E. coli* (70). However, roles for homologues have been identified and a role for YcaL may not be evident under the lab conditions tested.

Overall, our data reveals the power of high resolution TIS screens for understanding the contribution of genes to the overall fitness of a strain. We have revealed for the first time a direct link between iron availability and OM stability, potentially paving the way for future research into novel complementary therapeutic interventions to combat the rise of antimicrobial resistant infections.

## Materials and methods

### Strains and plasmids

**Table 1.** Strains and Plasmids used in this study.

| **Table 1.** Strains and Plasmids used in this study |  |  |
| --- | --- | --- |
| **Strains** | **Description** | **Reference** |
| S. Typhimurium SL1344 | Histidine auxotrophic variant S. Typhimurium | (13) |
| S. Typhimurium SL3261 | *aroA* deficient variant of SL1344 | (13) |
| *E. coli* DH5α | Cloning strain | NEB |
| **Plasmids** |  |  |
| pBAD18 | Arabinose inducible over expression vector | ATCC |
| pQE60*ndel* | T5 IPTG inducible vector | (71) |
| pQE60-*aroA*_his | *aroA* cloned into pQE60*ndel* encoding a C-terminal poly histidine tag. | This study |
| pQE60-*ycaL*-his | *ycaL* cloned into pQE60*ndel* encoding a C-terminal poly histidine tag. | This study |
| pQE60-*aroA*_ycaL-his | *aroA*-*ycaL* cloned into pQE60*ndel* encoding a C-terminal poly histidine tag. | This study |
| pBAD18-*aroA*-his | His-tagged AroA subcloned into pBAD18 using the *EcoRI* and *HindIII* sites. | This study |
| pBAD18-*ycaL*-his | His-tagged ycaL subcloned into pBAD18 using the *Sall* and *HindIII* sites. | This study |
| pBAD18-*aroA*-ycaL_his | AroA and His-tagged YcaL subcloned into pBAD18 using the *EcoRI* and *HindIII* sites. | This study |
| pLUX | Reporter vector containing bacterial luminescence gene operon. | (72) |
| pLUX-*pnrdA* | *Salmonella nrdA* promoter cloned between the *XhoI* and *BamHI* sites. | This study |
| pLUX-pnrdD | *Salmonella nrdD* promoter cloned between the *XhoI* and *BamHI* sites. | This study |
| pLUX-pnrdH | *Salmonella nrdH* promoter cloned between the *XhoI* and *BamHI* sites. | This study |
| pLUX-pgyrA | *Salmonella gyrA* promoter cloned between the *XhoI* and *BamHI* sites. | This study |

**Table 2.**
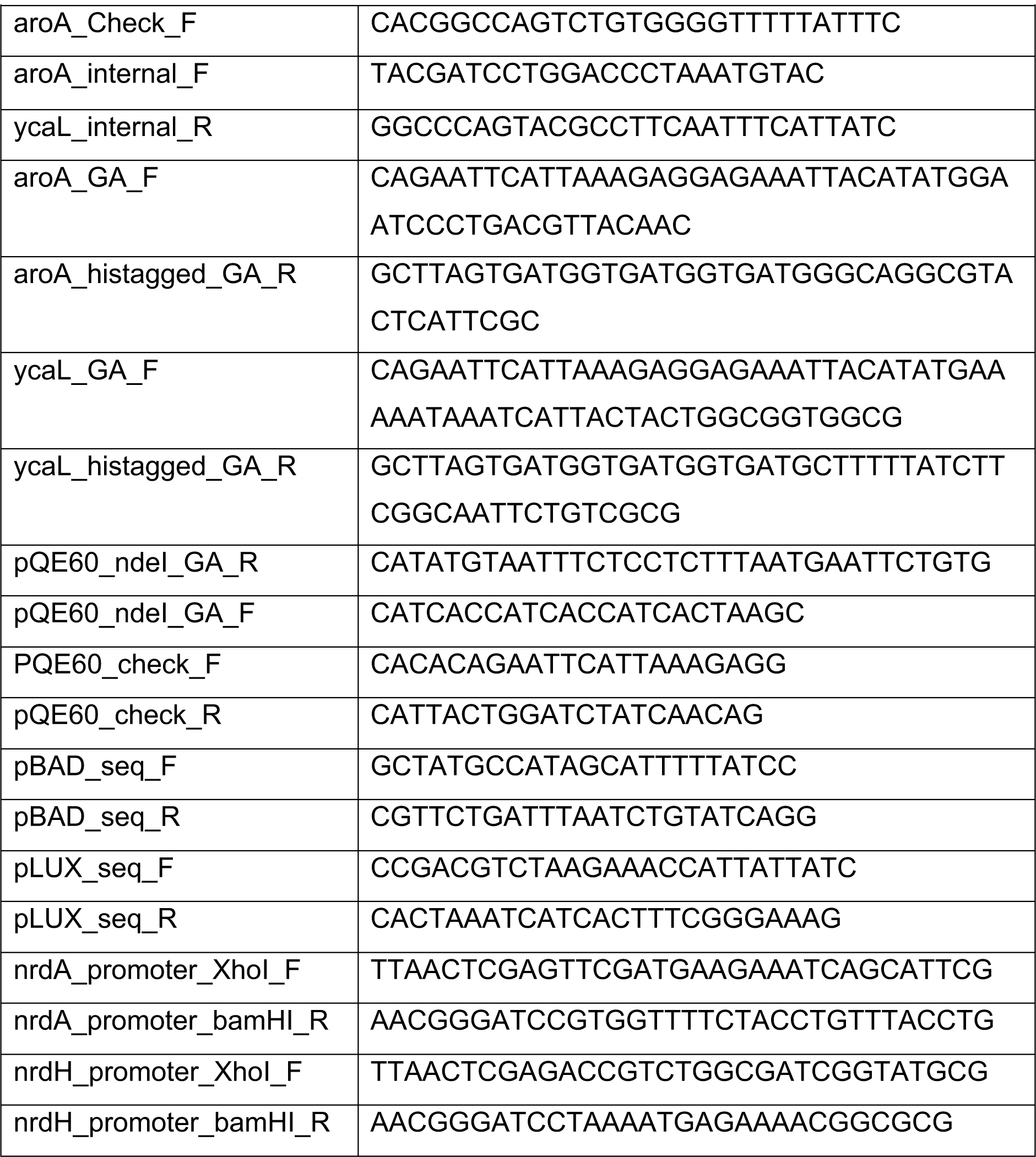
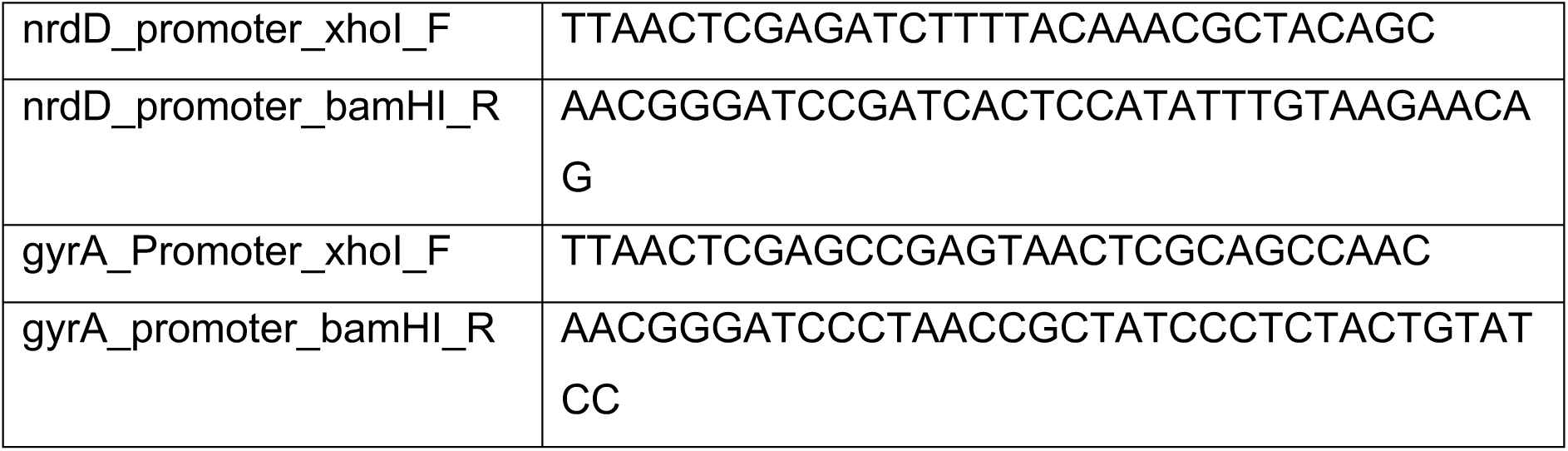
Primers used in this study.

### Media and growth conditions

All strains were grown in Luria-Bertani (LB) broth at 37°C with shaking unless stated otherwise. LB agar was made using 1.5% agar in LB. M9 minimal media was prepared using 5 x M9 salts (Sigma Aldrich) and supplemented with 200 mM MgSO4, 1 mM CaCl2, 30 μg/ml L-histidine and 0.4% (w/v) glucose. Growth curves were completed in 96 well plates and OD_600nm_ measured using a Clariostar plate reader.

### Molecular biology

pQE60*NdeI* vectors were generated using Gibson assembly (73). Briefly, the pQE60*NdeI* backbone was amplified by PCR using primer pair pQE60_ndeI_GA_F and pQE60_ndeI_GA_R. The aroA-his and aroA-ycaL-his inserts were also generated by PCR, with the primers including 20-30 bp homology with the target vector. Insert and vector PCR fragments were incubated with Gibson Assembly mastermix (NEB) for 1 h and 50°C. The assembled vectors were then transformed into NEB ultra competent *E. coli* DH5α cells using heat shock. Inserts were subcloned into pBAD18 using the *EcoRI* or *SalI* and *HindIII* restrictions sites. For cloning into pLUX, promoter regions from target genes were cloned into the *XhoI* and *BamHI* restrictions sites. All transformants were confirmed by PCR and Sanger sequencing.

### Protein sample preparation, SDS-PAGE and Western blotting

Approximately 10^9^ colony forming units (cfu) from overnight cultures were centrifuged and re-suspended in 2x Leammli sample buffer (Sigma). The samples were boiled and loaded onto 4-12% NuPage gels. After proteins were separated by SDS-PAGE electrophoresis, proteins were transferred to nitrocellulose membranes using a TurboBlot transfer system and blocked in 5% skim milk. Membranes were probed with either anti-RNAp antibodies (1:2,000) or anti-his antibodies (1:5,000). Secondary goat anti-mouse IgG IRDye 680RD antibodies were used to detect protein bands and the blots imaged using an Odyssey CLx machine.

### TraDIS library construction

Both the SL1344 and SL3261 transposon libraries were constructed using the following method. Electrocompetent cells were generated after growth at 37°C in 2xTY broth (12 g tryptone, 6 g yeast extract, 3.4 mM CaCL2 in 1 L) without salt until an optical density (OD_600_) of 0.6. Cells were pelleted at 5,000 x*g* and washed in iced cold sterile distilled water. The final cell pellet was resuspended in iced cold 10% glycerol (V/V). Aliquots of cells were mixed with EZ-Tn5^TM^ <KAN-2> transposome (Lucigen) and incubated on ice for 30 min. Cells were pulsed at 2500 V and pre-warmed SOC medium was immediately added to the sample for recovery. Cells were incubated at 37℃ for 2 h and transposon mutants selected for on LB agar plates supplemented with 50 µg/ml kanamycin. Plates were incubated at 37℃ for 18 h. Colonies were scraped from each plate and pooled to form the library, stored in a single falcon with 30% LB-glycerol.

### Transposon junction sequencing

Genomic DNA was extracted from each library using a RTP Stratec DNA extraction kit following the kit protocol. DNA was fragmented using a Covaris bioruptor to achieve a fragment size of ∼200 bp. The fragments containing the transposon-junctions were amplified by PCR and prepared for sequencing using the NEB Ultra I kit (New England Biolabs) following the manufacturer’s instructions. The final sample was quantified by qPCR using a Kapa Library Quantification kit for Illumina platforms. Samples were sequenced using an Illumina MiSeq, obtaining ∼5 million reads per sample. Data are deposited with the European Nucleotide Archive, accession number PRJEB65934.

### Sequencing data analysis

The data were processed first to remove any barcode using the fastx barcode splitter and trimmer tools from the Fastx toolkit. Sequence data with a correct barcode and transposon sequence, following trimming, were processed using Bio-Tradis and subsequently mapped to the SL1344 reference genome (accessions NC_016810.1; NC_017718.1; NC_017719.1; NC_017720.1). Comparative analyses between libraries was also done using the Bio-TraDIS tradis_comparisons.R script, using a minimum read count of 100 for comparison and 5% trim applied to both the 5´ and 3´ end of each gene. Data were inspected in the Artemis genome browser (74). The data can also be visualised in our online browser, available at https://tradis-vault.qfab.org/apollo/jbrowse.

### Growth in iron limiting conditions

LB medium was depleted of iron using the iron chelator 2-2bipyridyl. A concentration gradient was generated in a 96 well plate and bacteria were added to a starting OD_600nm_ of approximately 0.02. Growth (OD_600nm_) was measured every 30 minutes for 16 hours in a polarstar plate reader. For iron supplementation growth, iron was chelated using 600 µm of 2-2 bipyridyl and Fe^3+^ supplementation at 1 mM.

### Screening for outer membrane defects

Overnight cultures were diluted to approximately 10^9^ cfu/ml and serially diluted prior to spotting onto LB agar plates supplemented with either DMSO, 1 µM enterobactin, 200 µg/ml vancomycin, 2 mM EDTA or 1% bile salts (w/v). Plates were incubated overnight at 37°C for 16-18 h.

### Reporter screening

Testing promoter-reporters were completed under mid-exponential phase (MEP), low iron and anaerobic conditions as previously described (75). Bacteria were added to black walled 96 well plates and luminescence was measured in a Tecan plate reader and normalised to OD_600nm_. For reporter activity on agar, 100 µl of LB agar was added to black walled 96 well plates and 10^6^ cfu were inoculated into each well. OD_600nm_ and luminescence were measured in a Clariostar plate reader.

### LPS preparations

Approximately 10^9^ cfu of overnight cultures were pelleted and resuspended in Laemmli buffer. Cells were boiled prior to the addition of proteinase K and incubated for 1 h at 60°C. Proteinase K was heat inactivated prior to analysis by SDS-PAGE and silver staining.

## Supplementary Tables

**Table S1.** Transposon libraries constructed in *S.* Typhimurium strains.

| Strain | Transposon | Year | No. of mutants<sup>a</sup> | No. UIP<sup>b</sup> | Insertion density<sup>c</sup> | Reference |
| --- | --- | --- | --- | --- | --- | --- |
| SL1344 | Tn5/Mu | 2009 | ~10,000 | Not stated | N/A | (76) |
| SL1344 | Tn5 | 2016 | 4,381 | Not stated | N/A | (77) |
| SL1344 | Mu | 2016 | 4,975 | Not stated | N/A | (77) |
| SL3261 | Tn5 | 2013 | ~930,000 | 549,086 | 9 bp | (26) |
| D23580 | Tn5 | 2019 | Not stated | 797,000 | 6 bp | (33) |
| 14028s | Tn5 | 2017 | 325,000 | Not stated | N/A | (78) |

<sup>a</sup> The number of colonies used to construct the library.
<sup>b</sup> The number of unique insertion points (UIP) for the library if reported.
<sup>c</sup> Insertion density is the genome size divided by the number of insertions or mutants reported for that genome to give an approximation of 1 insertion every X bp as a measure of overall library density.

**Table S2.** Metrics for the TIS libraries generated in this study.

| **Sample** | **Replicate 1** | **Replicate 2** | **Combined** | **UIPs** |
| --- | --- | --- | --- | --- |
| SL1344 chromosome | 506,930 | 431,294 | 625,038 | 7.8 bp |
| SL1344 pCol1B9 | 16,882 | 13,834 | 20,545 | 4.2 bp |
| SL1344 pSLT | 18,120 | 15,723 | 20,017 | 4.7 bp |
| SL3261 chromosome | 534,096 | 440,335 | 632,132 | 7.7 bp |
| SL3261 pCol1B9 | 18,250 | 17,772 | 23,431 | 3.7 bp |
| SL3261 pSLT | 15,496 | 16,371 | 20,194 | 4.6 bp |

## Supplementary Files

**Table S3. *S.* Typhimurium essential genes comparisons**

**Table S4. Mutant fitness comparisons.**

## Supplementary Figures

**Figure. S1.**
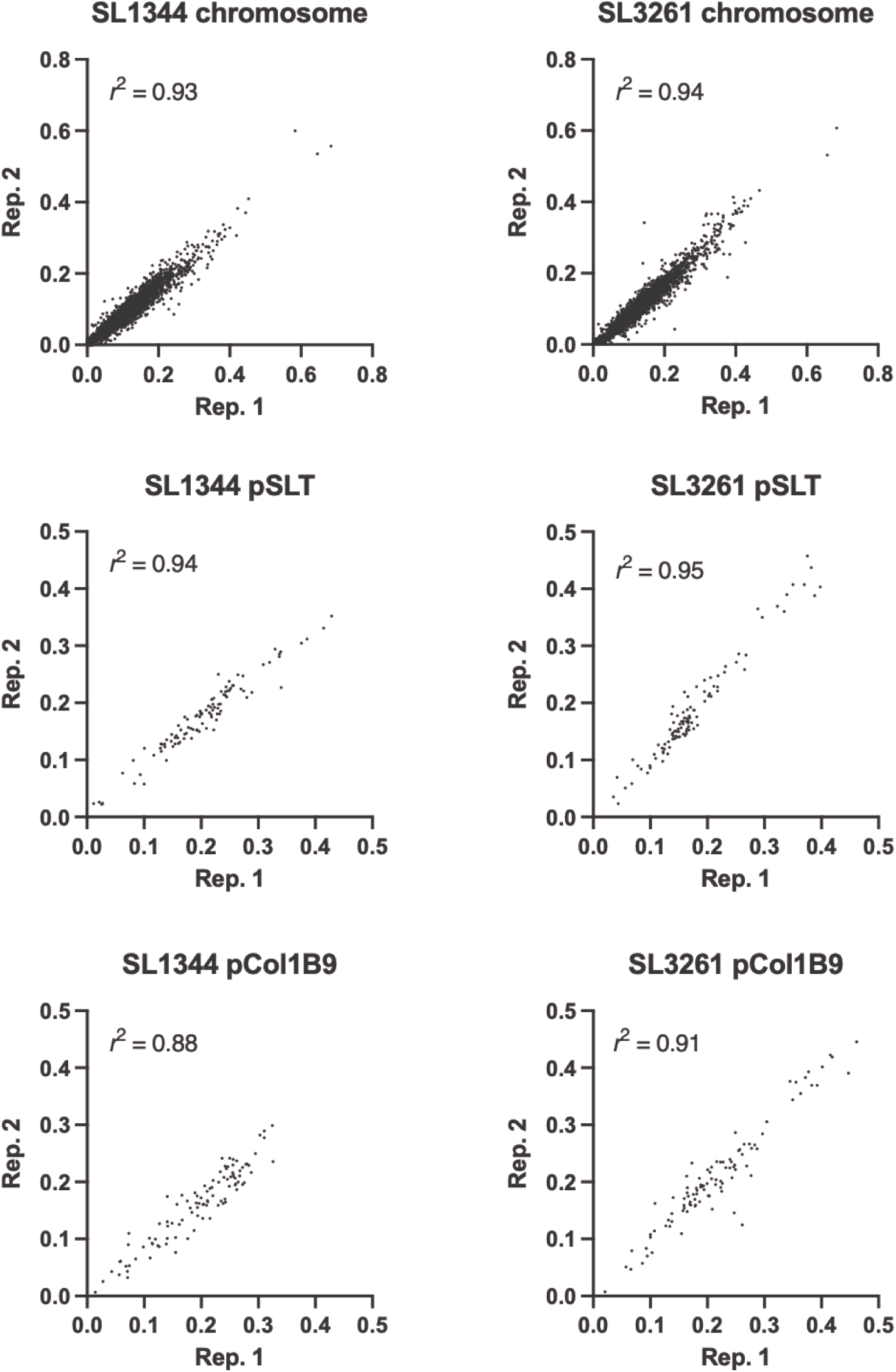
Comparison of technical replicates. Scatter graphs showing the insertion index scores of each technical replicate for SL1344 (left) and SL3261 (right) chromosome (top), pSLT (middle) and pCOL1B9 (bottom) plasmids.

**Figure S2.**
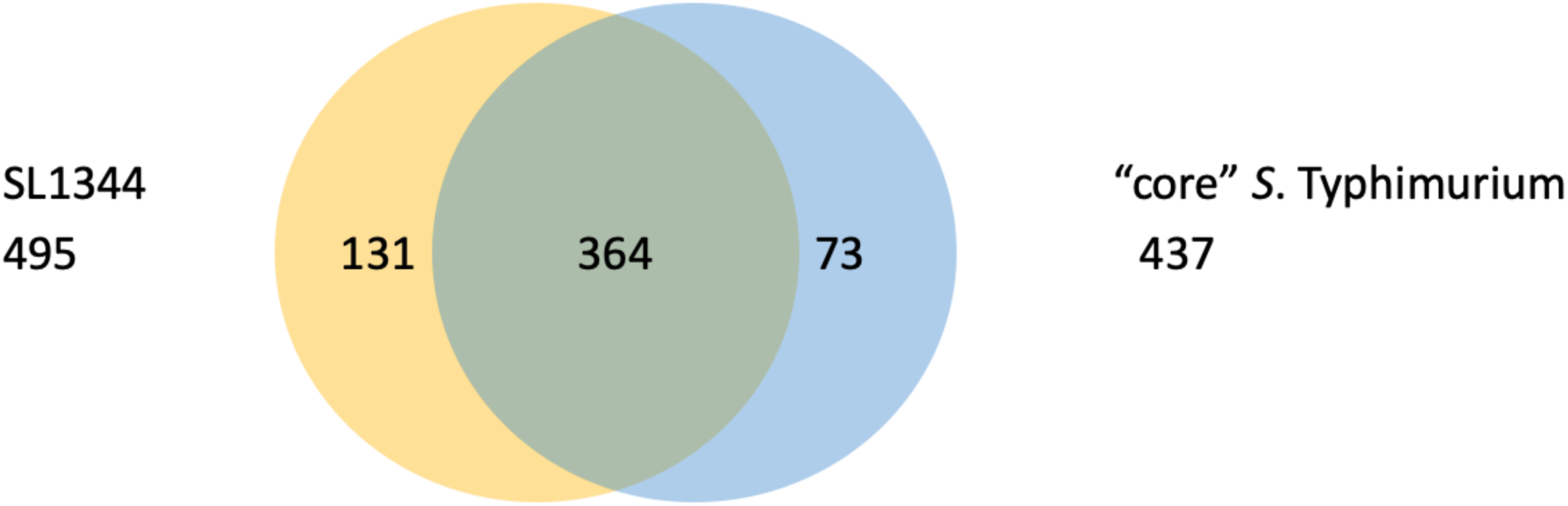
S. Typhimurium essential gene comparisons. Venn diagram over essential SL1344 genes identified in this stufy conpared to a compendium of S. Typhimurium essential genes collated from (33).

**Figure S3.**
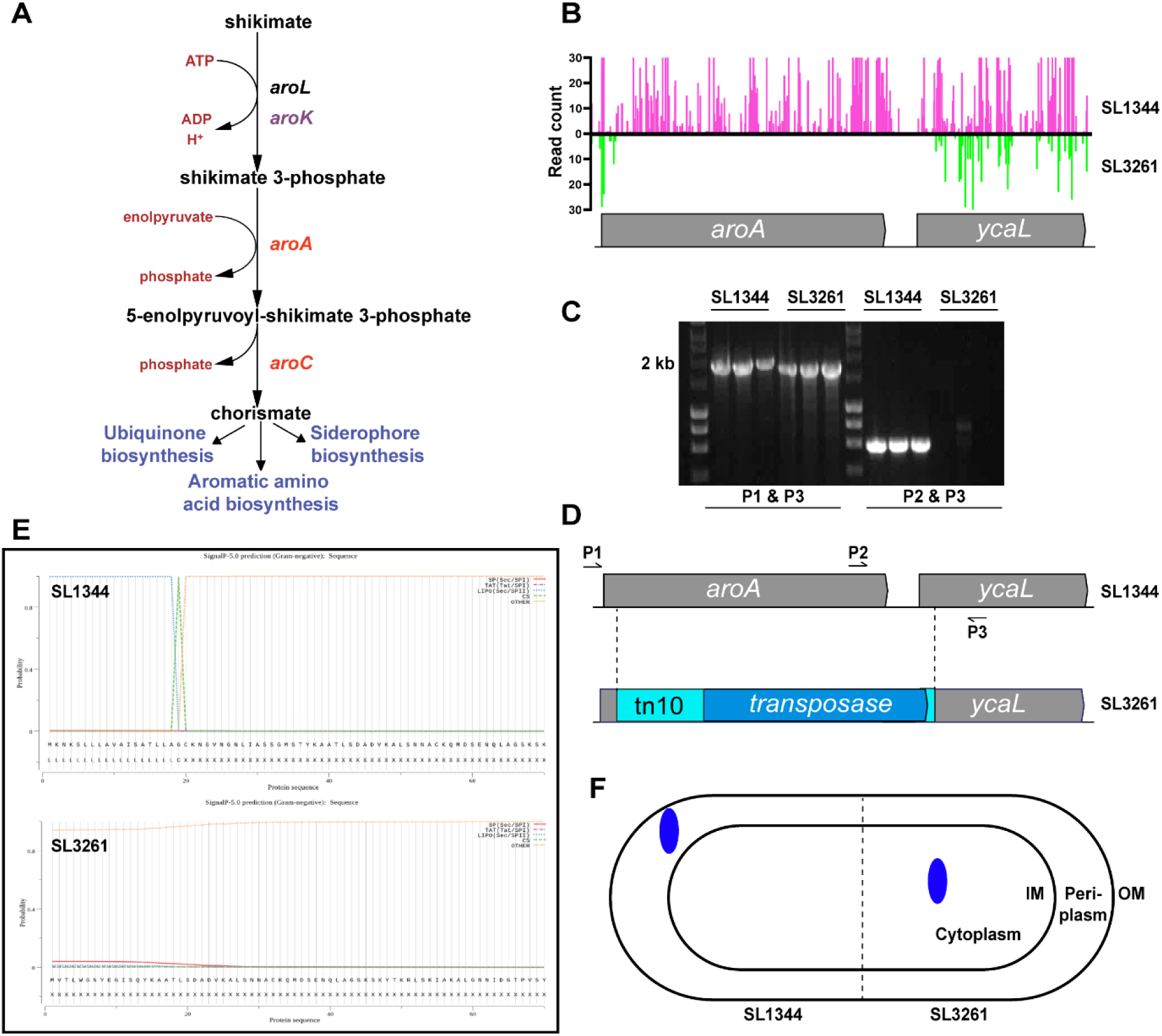
AroA-YcaL loci in *S*. Typhimurium. (A) Shikimate pathway in *S*. Typhimurium. (B) Insertion plot files for the *aroA-ycaL* loci in S. Typhimurium SL1344 (pink) and SL3261 (green) TIS libraries. (C) PCR amplification of the *aroA-ycaL* loci in SL1344 and SL3261. (D) Schematic of the genomic scar left by the tn10 transposon in SL3261. (E) SignalP plots for the YcaL amino acid sequence in SL1344 (top) and SL3261 (bottom). (F) Schematic of predicted YcaL protein localisation in both SL1344 and SL3261.

**Figure S4.**
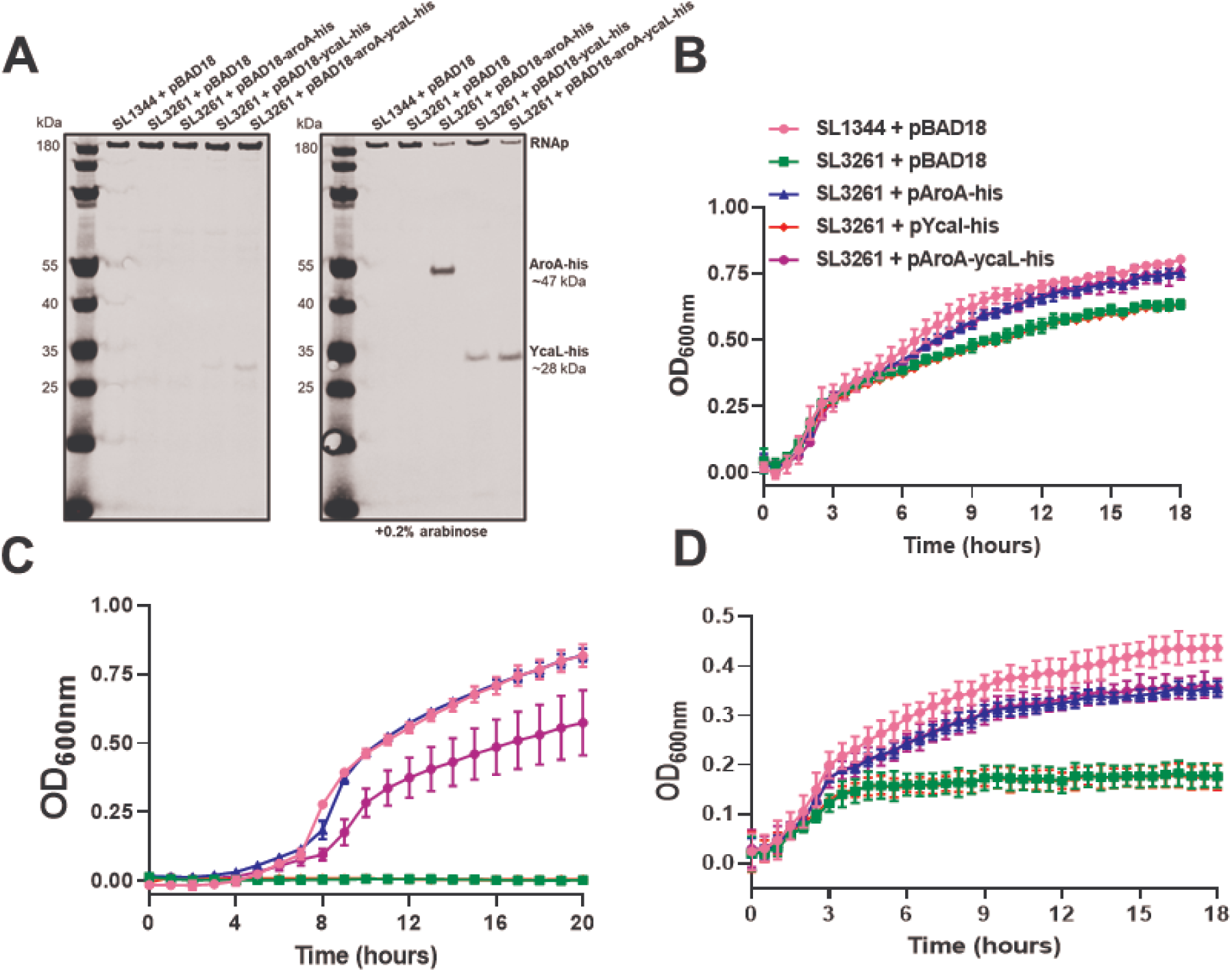
Complementation plasmids in SL3261. (A) Western immunoblot using anti-poly Histidine tag and anti-RNA polymerase antibodies. Whole cell protein samples were separated on 6-12 NuPAGE gels prior to Western blotting. Images were detected using fluorescent secondary antibodies. Growth of strains in (B) LB medium, (C) M9 minimal medium supplemented with 0.4% glucose and (D) LB medium with 600 µM 2-2 bipyridyl.

**Figure S5.**
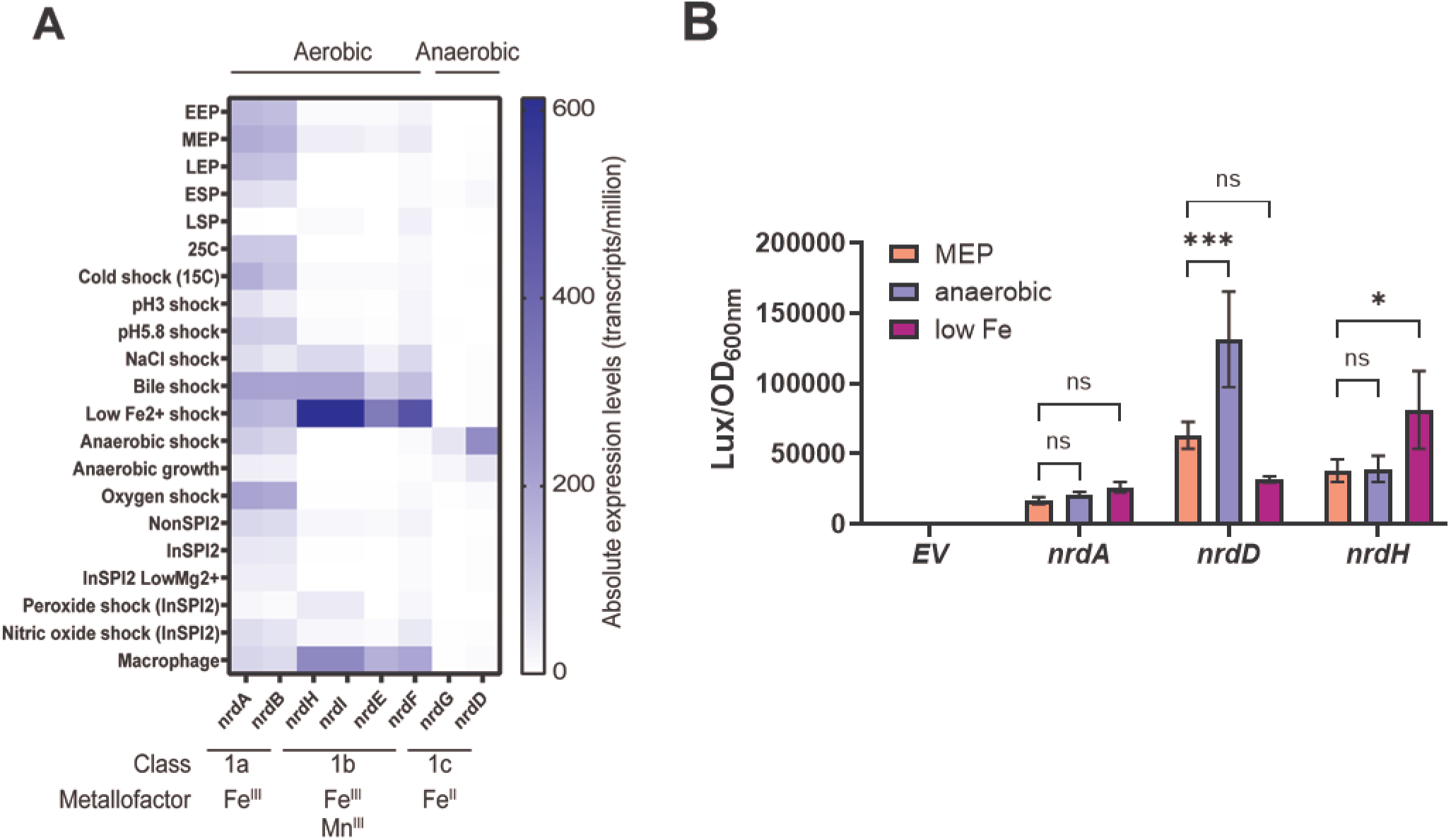
Differential regulation of S. Typhimurium RNR genes. (A) Absolute expression levels of NrdAB, NrdHIEF and NrdGD under multiple conditions determined by RNA-seq (49). (B) Nrd-luminescence reporter construct activity in mid exponential phase (MEP), anaerobic and low iron conditions.

**Figure S6.**
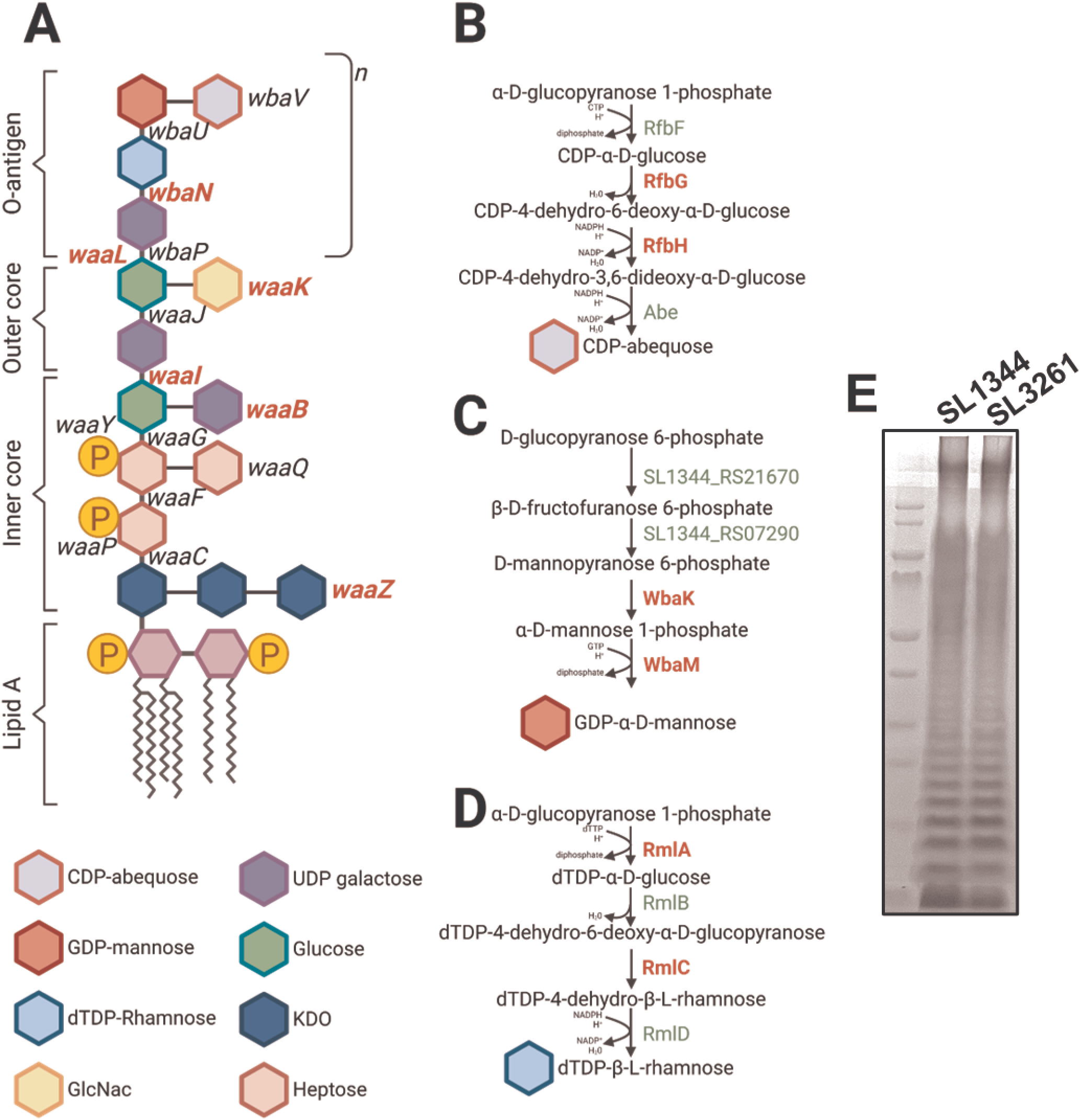
Mutant fitness of genes involved in LPS biosynthesis. Enzymes highlighted in red were identified in this study as having transposon mutants that were more fit in SL3261. (A) LPS structure and enzymes involved in the addition of each sugar onto the LPS structure. Sugar-nucleotide biosynthetic pathways for o-antigen sugar units (B) CDP-abequose, (C) GDP-α-mannose and (D) dTDP-β-rhamnose. (E) LPS profiles of SL1344 and SL3261.

## Notes

### Competing Interest Statement

The authors have declared no competing interest.

